# Aurora B-mediated exclusion of HP1a from late-segregating chromatin prevents formation of micronuclei

**DOI:** 10.1101/268912

**Authors:** Brandt Warecki, William Sullivan

## Abstract

Late-segregating acentric chromosomes pose a serious risk to genomic integrity when they are excluded from dividing daughter nuclei and form damage-prone micronuclei. Insight into the cellular mechanisms that prevent the formation of micronuclei from acentrics come from studies demonstrating that acentrics reincorporate into daughter telophase nuclei by passing through Aurora B kinase-dependent channels in the nuclear envelope of *Drosophila* neuroblasts. Here, we uncover a mechanism of nuclear envelope channel formation in which localized concentrations of Aurora B preferentially phosphorylate H3(S10) on heterochromatic acentrics and their associated DNA tethers. This phosphorylation event prevents HP1a from associating with heterochromatin and results in localized inhibition of nuclear envelope reassembly on endonuclease- and X-irradiation-induced acentrics and the main daughter nuclei at the sites of acentric entry to promote the formation of channels. Finally, we find that HP1a also specifies initiation sites of nuclear envelope reassembly on undamaged chromatin. Taken together, these results demonstrate that Aurora B-mediated regulation of HP1a-chromatin interactions plays a key role maintaining genome integrity by locally preventing nuclear envelope assembly and facilitating incorporation of late-segregating acentrics into daughter nuclei.

## INTRODUCTION

Eukaryotic cells have evolved sophisticated mechanisms that maintain genome integrity. Checkpoints halt cell cycle progression in response to damaged DNA to allow for repair or elimination of compromised cells [1]. For example, the G1-S and the G2-M checkpoints prevent entry into S-phase and mitosis respectively when DNA is damaged [2]. Additional checkpoints at the metaphase-anaphase transition delay progression into anaphase if DNA is damaged once a cell commits to mitosis [3–4]. Despite these checkpoints, cells sometimes enter anaphase with damaged DNA. Unrepaired double-stranded DNA breaks are particularly problematic, as they result in chromosome fragments lacking either a telomere or centromere [5]. The latter, called acentrics, are unable to form traditional microtubule-kinetochore attachments and are therefore expected to fail to segregate properly and to be excluded from the nascent daughter nuclei, leading to the formation of micronuclei [6–8]. Historically, micronuclei have served as a biomarker for cancerous tissue [9–10], and recent studies reveal that micronuclei drive genomic instability either through their loss during subsequent cell divisions or through chromothripsis, the dramatic shattering and rearrangement of micronuclear DNA that is then incorporated into the genome [11–14].

While the formation of micronuclei from lagging chromosomes has been widely documented, surprisingly, in some instances, discrete lagging chromosomes avoid micronuclei formation by rejoining daughter nuclei before mitosis is completed. For example, in human colorectal cancer cells, a proportion of nocodazole-induced lagging whole chromosomes that would otherwise form micronuclei instead reincorporate into the main daughter nuclei in late anaphase [15]. In fission yeast, certain lagging chromatids that stay distinct from the main segregating chromosomes during anaphase eventually reunite with daughter nuclei in telophase [16–17]. In addition, in *Drosophila* neuroblast and papillar divisions, late-segregating acentric fragments induced by endonuclease activity or irradiation successfully rejoin daughter nuclei in late telophase [18–19]. Therefore, the fate of lagging acentric chromosomes is an important but underexplored area of cell biology. Here, we specifically examine the mechanisms that facilitate incorporation of late-segregating acentric chromosomes into daughter nuclei, avoiding micronuclei formation.

In *Drosophila*, acentric behavior has been studied using transgenic flies containing a heat-shock inducible I-CreI endonuclease [18–24], which targets rDNA near the base of the X chromosome and creates double-stranded DNA breaks [20, 25–27]. I-CreI-mediated DNA breaks result in γH2Av foci that persist through mitosis and chromosome fragments that do not recruit canonical centromere components and thus are considered acentrics [18]. Even though I-CreI-induced acentrics initially lag on the metaphase plate while undamaged chromosomes segregate, acentrics ultimately undergo delayed but successful segregation [18]. Acentric segregation is achieved through protein-coated DNA tethers connecting acentrics to their centric partners and microtubule bundles that encompass acentrics, enabling their poleward movement [24]. The histone-based DNA tether is associated with Polo, BubR1, and the chromosome passenger proteins Aurora B and INCENP [18].

Because lagging and acentric chromosome segregation is significantly delayed, occurring late in anaphase, lagging and acentric chromosomes often remain distinct from the main mass of chromosomes after nuclear envelope reassembly has been initiated [22, 28–30]. Despite the presence of the nascent nuclear envelope surrounding the main nuclear mass, in *Drosophila* neuroblasts, lagging acentrics are not “locked out” of daughter nuclei and do not form micronuclei. Rather, the late-segregating acentrics bypass the nuclear envelope barrier and enter telophase nuclei through channels in the nuclear envelope that are formed by highly localized delays in the completion of nuclear envelope reassembly [22]. Nuclear envelope channel formation is dependent upon the Aurora B kinase activity that is associated with the acentric and DNA tether. When Aurora B activity is reduced, acentrics are unable to enter daughter nuclei and instead form lamin-coated micronuclei. Presumably, the pool of Aurora B responsible for channel formation comes from Aurora B persisting on the DNA tethers and acentrics, as channel formation is not observed in divisions which lack both acentrics and their associated Aurora B-coated tethers [22].

The highly localized delays in nuclear envelope reassembly resulting in nuclear envelope channels suggest localized inhibition of important steps in nuclear envelope reassembly. Key events in nuclear envelope reassembly include reformation of nuclear pore complexes, reestablishment of important connections between chromatin and inner nuclear membrane proteins that are disrupted in early mitosis, and fusion of nuclear envelope membrane microdomains [31–36]. Additionally, the nuclear lamina reassembles once nuclear pore complexes and inner nuclear membrane proteins are recruited to daughter nuclei [37–40]

Regulation of these nuclear envelope reassembly events is achieved through the global activity of mitotic kinases, among which Aurora B is a known negative regulator [22, 30, 41]. One mechanism by which Aurora B activity may inhibit nuclear envelope reassembly is through disrupting chromatin interactions with the heterochromatin component HP1α/HP1a [42]. In interphase, HP1α/HP1a interacts both with methylated histone H3 and with nuclear envelope components [43–46]. As cells enter mitosis, Aurora B-mediated phosphorylation of H3(S10) acts as a switch to remove HP1α/HP1a from chromosomes [47–49]. During anaphase, when Aurora B relocalizes to the spindle midzone and H3(S10) phosphate groups are removed [50], HP1α/HP1a is reloaded onto segregating chromosomes and subsequently reestablishes connections with nuclear envelope-associated components [51–52], which is a possible early step in reformation of the nuclear envelope [45].

Understanding the mechanisms by which Aurora B kinase activity locally alters the events of nuclear envelope reassembly to mediate channel formation is of particular interest. Specifically, understanding the pathway through which Aurora B acts to form channels and allow incorporation of late-segregating acentrics into daughter nuclei would reveal new mechanisms by which Aurora B prevents micronuclei formation and maintains genome integrity. In addition, studying the mechanisms by which nuclear envelope channel formation is regulated may provide a system for understanding mechanisms that regulate global nuclear envelope reassembly in wild-type divisions. Based on the data presented here, in which we generate acentrics using both the I-CreI endonuclease and X-irradiation, we propose a model for nuclear envelope channel formation in which highly localized concentrations of Aurora B kinase phosphorylate H3(S10) specifically on acentrics and their associated tethers. This prevents local heterochromatic recruitment of HP1a and subsequent recruitment of lamin, nuclear pore complexes, and nuclear envelope membrane at the sites where acentrics rejoin daughter nuclei (Figure 1).

**Figure 1.**
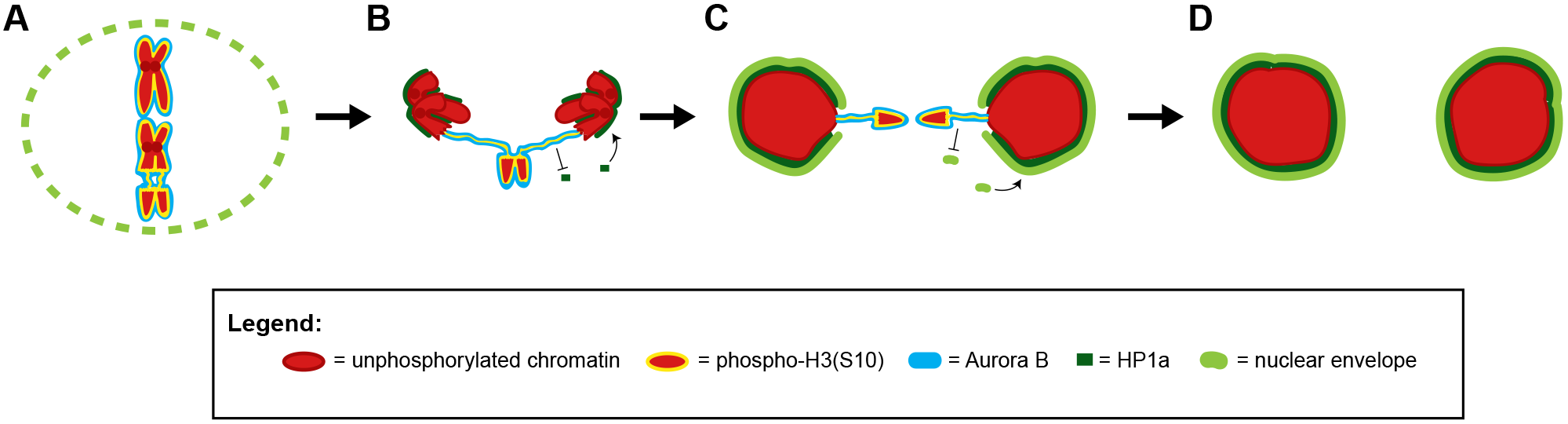
Model for Aurora B-mediated nuclear envelope channel formation. (A) During metaphase, Aurora B (blue) phosphorylates H3(S10) (yellow) on chromosomes (red), acentrics are pushed to the edge of the metaphase plate, and the nuclear envelope (lime green) is partially disassembled. (B) During anaphase, Aurora B, a component of the chromosome passenger complex, is removed from the main chromosomes and relocalizes to the spindle midzone. Phospho-H3(S10) marks on the main nuclei are removed, and HP1 (dark green) is recruited to the main nuclei. However, persistent Aurora B on the acentric and tether continues to phosphorylate H3(S10) and inhibits HP1a recruitment to the acentric/tether. (C) During telophase, nuclear envelope components reform connections with chromatin through HP1a and nuclear envelope reassembly proceeds. However, the exclusion of HP1a from Aurora B-coated tether/acentrics prevents accumulation of nuclear envelope components on the tether/acentrics and at the site where the tether contacts the main nucleus, leading to local delays in nuclear envelope reassembly and the formation of nuclear envelope channels. (D) Successful incorporation of late-segregating acentrics into telophase nuclei through nuclear envelope channels results in euploid daughter cells.

## RESULTS

### Aurora B kinase preferentially phosphorylates H3(S10) on acentrics and on chromatin near sites of channel formation

Aurora B kinase, a component of the chromosome passenger complex, initially localizes to and modifies chromosomes in early mitosis and then localizes to the spindle midzone during anaphase [50]. Previous studies demonstrated that Aurora B kinase persists on DNA tethers stretching from lagging acentrics to the newly formed daughter nuclei despite its removal from the main mass of segregating chromosomes [18, 22]. This highly localized Aurora B activity mediates nuclear envelope channel formation to allow acentric entry into telophase nuclei [22]. Thus, we tested if the Aurora B localized on the acentric and associated tether maintains the acentric chromatin and chromatin at the site where the acentric enters the main nucleus in a mitotic state that would locally prohibit nuclear envelope formation.

Aurora B modifies mitotic chromatin through phosphorylation of histone H3 on serine 10 in a spatiotemporal manner [53–55]. Therefore, we tested whether Aurora B-dependent phosphorylation of H3(S10) persists on acentrics, tethers, and channel sites despite this mark having been removed from the main nuclei. We fixed I-CreI-expressing mitotic 3^rd^ instar larval neuroblasts and stained with an antibody that specifically recognizes phospho-H3(S10) (Figure 2). In neuroblasts fixed in metaphase, we observed phospho-H3(S10) present along the length of all chromosomes, consistent with data from [56] (Figure 2A, left panel). In anaphase neuroblasts, we observed a weak phospho-H3(S10) signal on segregating intact chromosomes. In contrast, late-segregating acentrics that remained at or had just segregated from the metaphase plate (Figure 2A, middle panels, red arrows) exhibited a strong phospho-H3(S10) signal (Figure 2A, middle panels, cyan arrows). In neuroblasts fixed in telophase, the stage at which acentrics begin to rejoin daughter nuclei, we observed strong phospho-H3(S10) signal on acentrics. The phospho-H3(S10) signal abruptly ended at the point of contact between the acentric and daughter nucleus (Figure 2A, right panel).

**Figure 2.**
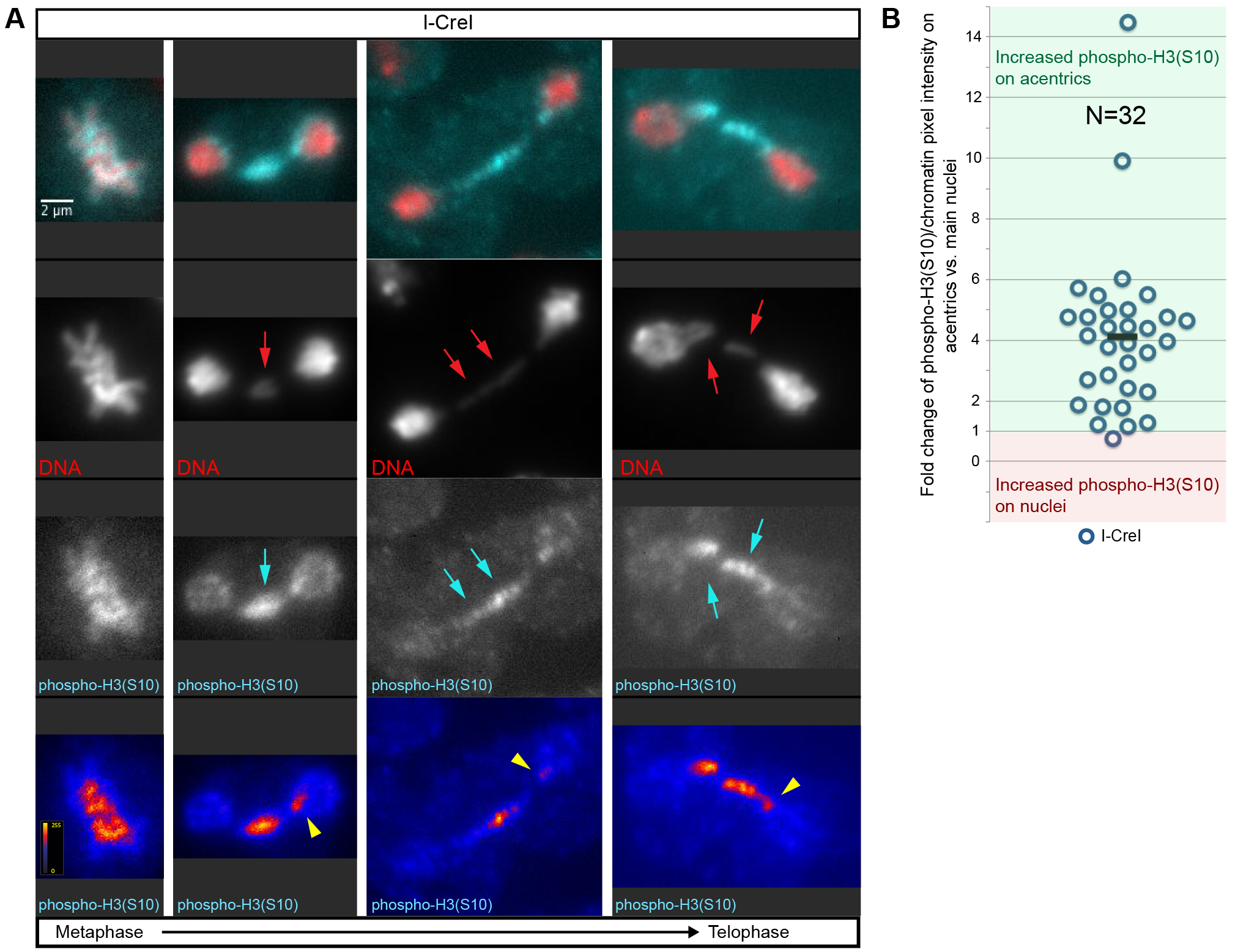
H3(S10) is preferentially phosphorylated on acentrics and tethers. (A) Fixed mitotic neuroblasts expressing I-CreI in metaphase (left panel), anaphase (middle panels), and telophase (right panel) (red = DNA; blue = phospho-H3(S10)). In metaphase, phospho-H3(S10) was detected coating all chromosomes. During anaphase, a strong phospho-H3(S10) signal (cyan arrows) was detected on acentrics (red arrows), but diminished on the main segregating chromosome complements. By telophase, little to no phospho-H3(S10) was detected on the main nuclei, yet late-segregating acentrics (red arrows) were associated with increased phosphorylation of H3(S10) (cyan arrows). Intriguingly, 47% (N=19) of the anaphase neuroblast nuclei possessed phospho-H3(S10) “hotspots” (yellow arrowheads) at the site of acentric entry. (B) Graphical comparison of fold increases in the average phopsho-H3(S10)/DNA pixel intensity ratio for the areas of the acentrics vs. the areas of the main nuclei for I-CreI-expressing neuroblasts. Each circle represents one anaphase/telophase cell. Values above 1 (green box) indicate increased phospho-H3(S10)/DNA pixel intensity ratio on acentrics. Values below 1 (red box) indicate increased phospho-H3(S10)/DNA pixel intensity ratio on main nuclei. 31/32 l I-CreI neuroblasts exhibited increased phospho-H3(S10)/DNA ratios on acentrics compared to main nuclei (Black bar: mean fold change = 4.13; SD = 2.65; N = 32). Scale bars are 2 μm. See also Figure S1.

To quantify these observations, we measured the ratio of signal intensities of phospho-H3(S10)/DNA on main intact nuclei and acentrics and then analyzed the fold change of this ratio for each fixed neuroblast division imaged (Figure 2B). Strikingly, for 31 out of the 32 neuroblast divisions scored, we observed an increase in phospho-H3(S10)/DNA intensity on acentrics compared to main intact nuclei (mean fold change = 4.13 +/− 2.65; N=32), consistent with previous reports [54–55].

Intriguingly, in a large proportion (47%; N=19) of anaphase- or telophase-fixed neuroblast divisions in which at least one acentric remained distinct from the main nuclei, we clearly detected localized “hotspots” of strong phospho-H3(S10) intensity on one of the newly formed main nuclei at presumptive sites of acentric entry. These hotspots correspond to the location where tethers contact the main nuclei and nuclear envelope channel formation is generally observed (Figure 2A, yellow arrowheads). Taken together, these data demonstrate that acentrics, tethers, and the chromatin at sites of channel formation remain preferentially phosphorylated on H3(S10) even though phosphorylation of H3(S10) is broadly reduced on the chromatin in the newly formed telophase nuclei.

To determine whether Aurora B kinase activity is responsible for the observed preferential phosphorylation of H3(S10) on acentrics and tethers, we compared the fold changes of phospho-H3(S10)/DNA intensity on acentrics to main intact nuclei between neuroblasts with normal and RNAi-reduced levels of Aurora B. In I-CreI-expressing neuroblasts fixed in anaphase and telophase (Figure S1A), we observed an average fold change of phospho-H3(S10)/DNA signal on acentrics to main nuclei of 6.45 +/− 6.56 (N=23) with 78% of divisions showing a greater than 2-fold increase of phospho-H3(S10)/DNA intensity on acentrics compared to the main nuclei (and 57% of divisions showing a greater than 4-fold increase of phospho-H3(10)/DNA intensity on acentrics compared to the main nuclei) (Figure S1C).

In contrast, upon reduction of Aurora B (Figure S1B), we observed a decreased average fold change of phospho-H3(S10)/DNA signal on acentrics compared to main nuclei of 2.34 +/− 2.15 (N=20) with only 40% of divisions showing a greater than 2-fold increase of phospho-H3(S10)/DNA intensity on acentrics to main nuclei (compare to 78% of acentrics in wild-type conditions, significant by chi-square test *p*=0.0105) (and only 20% of divisions showing a greater than 4-fold increase of phospho-H3(S10)/DNA intensity on acentrics compared to the main nuclei) (Figure S1C). These findings indicate that Aurora B kinase is responsible for the observed persistent phosphorylation of H3(S10) on acentrics during anaphase and telophase.

### Aurora B kinase activity blocks HP1a association on late-segregating acentrics

I-CreI expression results in acentrics containing a significant proportion of heterochromatin, which would be expected to recruit heterochromatic proteins to the acentric fragments. Since phosphorylation of H3(S10) by Aurora B kinase is known to prevent H3 interaction with the heterochromatin component HP1α (the mammalian ortholog of HP1a) [47–48], we hypothesized that the observed increase in phosphorylation of H3(S10) on acentrics with respect to the main nuclei would lead to an Aurora B-dependent preferential exclusion of HP1a on late-segregating acentrics despite HP1a recruitment to the main intact chromosomes. To test this hypothesis, we performed live imaging of dividing neuroblasts expressing I-CreI, H2Av-RFP, and GFP-HP1a in control conditions (Dimethyl sulfoxide (DMSO)-treated neuroblasts) and conditions in which Aurora B kinase activity was partially inhibited through introduction of the Aurora B-specific small molecule inhibitor Binucleine-2 [57] (Figure 3).

**Figure 3.**
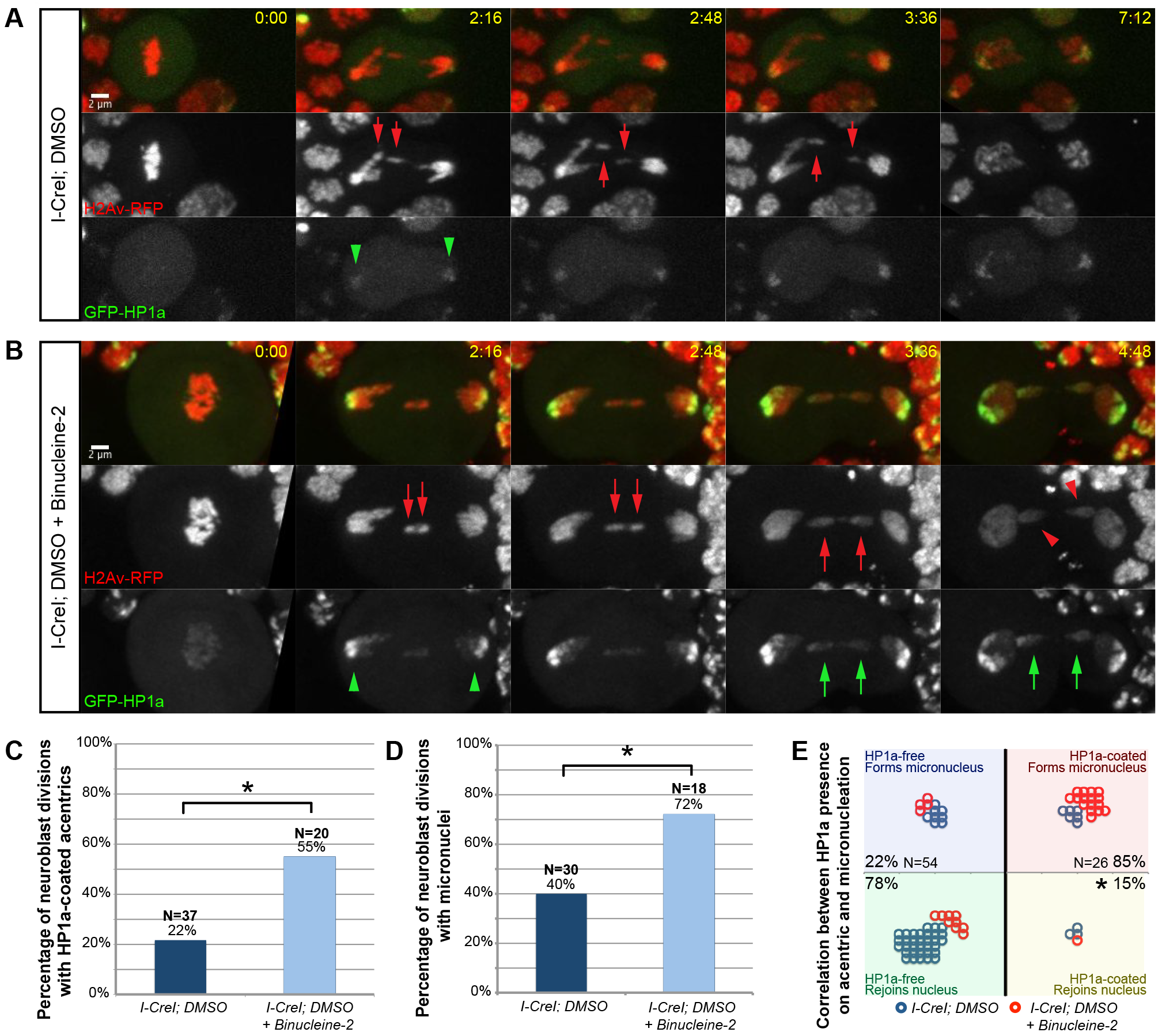
Aurora B kinase activity blocks HP1a association with late-segregating acentrics. (A) Stills from a time-lapse movie of a mitotic neuroblast expressing I-CreI, H2Av-RFP (red), and GFP-HP1a (green) treated with DMSO (control) (see Movie S1). 2:16 after anaphase onset, GFP-HP1a was detected on the centric heterochromatic region of segregating intact chromosomes (green arrowheads). However, throughout anaphase, no GFP-HP1a was detected on the pericentric heterochromatin-containing acentrics (red arrows), which separated and rejoined daughter nuclei. (B) Stills from a time-lapse movie of a mitotic neuroblast expressing I-CreI, H2Av-RFP (red), and GFP-HP1a (green) treated with the Aurora B inhibitor Binucleine-2 dissolved in DMSO (see Movie S2). At 2:16 after anaphase onset, GFP-HP1a was strongly detected on the centric heterochromatin of segregating intact chromosomes (green arrowheads). In addition, by 3:36 post-anaphase, increased GFP-HP1a was observed on acentric chromosomes (green arrows), which segregated but failed to rejoin daughter nuclei, instead forming GFP-HP1a-coated micronuclei (red arrowheads). (C) Comparison of the percentage of divisions in which GFP-HP1a was detected on acentrics in DMSO-treated (left) and DMSO + Binucleine-2-treated (right) I-CreI-expressing neuroblasts: 22% (N=37) and 55% (N=20) respectively. Asterisk indicates statistical significance (p=0.01). (D) Comparison of the percentage of divisions in which acentrics formed micronuclei in DMSO-treated (left) and DMSO + Binucleine-2-treated (right) I-CreI-expressing neuroblasts: 40% (N=30) and 72% (N=18) respectively. Asterisk indicates statistical significance (p=0.003). (E) Graph depicting a strong correlation between the lack of HP1a on the acentric and the ability of the acentric to rejoin the daughter nucleus. Each circle represents one acentric: blue and red circles depict single acentrics derived from DMSO and DMSO + Binucleine-2 treated I-CreI-expressing neuroblasts respectively. 78% of acentrics with no clear GFP-HP1a association (N=54) rejoined daughter nuclei, while 85% of acentrics with clear GFP-HP1a association (N=26) formed micronuclei. Asterisk indicates statistical significance (p=1.2 × 10^−7^). Time is written as min:sec after anaphase onset. Scale bars are 2 μm.

In control DMSO-treated mitotic neuroblasts (Figure 3A, see Movie S1), we observed the following patterns of HP1a association: in metaphase, little or no HP1a was detected on the chromosomes; and in anaphase, a strong HP1a signal was detected on the main segregating chromosomes (Figure 3A, green arrowheads). These results are consistent with previous data showing that a large proportion of HP1α/HP1a dissociates from chromosomes in early mitosis and re-associates with segregating chromosomes in anaphase [47–49, 52]. Interestingly, despite the anaphase recruitment of HP1a on the main intact chromosomes, little or no HP1a was detected on late-segregating acentrics (Figure 3A, red arrows). Acentrics segregated normally and rejoined daughter nuclei in telophase. In total, we only clearly detected HP1a on acentrics in about 22% of neuroblast divisions we imaged (N=37) (Figure 3C) and observed micronucleation in 40% of divisions (N=30) (Figure 3D).

In mitotic neuroblasts that were treated with the Aurora B inhibitor Binucleine-2 dissolved in DMSO (Figure 3B, see Movie S2), we observed similar patterns of HP1a association on the main intact nuclei: little or no HP1a on metaphase chromosomes followed by anaphase recruitment of HP1a on the main segregating chromosomes (Figure 3B, green arrowheads). However, in contrast to DMSO-treated neuroblasts, in which HP1a was not detected on late-segregating acentrics, we observed strong HP1a association on anaphase acentrics in neuroblasts treated with DMSO + Binucleine-2 (Figure 3B, green arrows). Acentrics failed to rejoin daughter nuclei, forming micronuclei (Figure 3B, red arrowheads). In all the DMSO + Binucleine-2-treated neuroblasts imaged, clear HP1a association was observed 55% of the time (N=20) (Figure 3C) (compare to 22% of the time in control divisions, a statistically significant difference) and micronucleation 72% of the time (N=18) (Figure 3D) (a statistically significant increase compared to the 40% micronucleation observed in control divisions), indicating that Aurora B kinase activity preferentially excludes HP1a from late-segregating acentrics.

We next determined whether HP1a association with acentrics was correlated with micronucleation. We scored individual acentrics for the presence of HP1a and whether the acentric formed a micronucleus (Figure 3E). We grouped the scored acentrics into four categories: 1) acentrics that were HP1a-free and formed micronuclei (Figure 3E, blue box); 2) acentrics that were HP1a-free and rejoined daughter nuclei (Figure 3E, green box); 3) acentrics that were HP1a-coated and formed micronuclei (Figure 3E, red box); and 4) acentrics that were HP1a-coated and rejoined daughter nuclei (Figure 3E, yellow box). Overall, we found that 78% of acentrics that were HP1a-free (N=54) rejoined daughter nuclei, and 85% of acentrics that were HP1a-coated (N=26) formed micronuclei. Thus, the absence or presence of HP1a is a strong predictor of the fate of the I-CreI-induced acentric: entering the daughter nucleus or forming a micronucleus.

### Aurora B kinase activity promotes nuclear envelope channel formation through HP1a exclusion from acentrics and tethers

Due to the correlation of HP1a-acentric association and the formation of micronuclei as well as the ability of HP1α/HP1a to interact with the nuclear envelope [43, 45], we tested whether Aurora B-mediated HP1a exclusion from acentrics might be a key factor in channel formation through localized inhibition of nuclear envelope reassembly. We performed live imaging of mitotic neuroblasts expressing I-CreI, H2Av-RFP, and the nuclear envelope marker Lamin-GFP and asked whether depletion of HP1a rescued the ability to form nuclear envelope channels when Aurora B was inhibited. The rationale is that while chromatin containing H3-HP1a can be a strong promoter of nuclear envelope assembly, chromatin containing H3 by itself is only a neutral substrate for assembly, and thus prevention of H3-HP1a formation on the acentric and tether, either through Aurora B-mediated phosphorylation of H3(S10) or through depletion of HP1a, would be conducive to channel formation (Figure S2).

HP1α/HP1a is essential: homozygous null mutants result in embryonic lethality in *Drosophila* [58]. Therefore, to address the role of HP1a in nuclear envelope channel formation, we made use of the transgenic UAS/Gal4 system (for review, see [59]) to deplete HP1a through RNA interference (RNAi). In our setup, we used elav-Gal4 to drive expression of UAS-HP1a-dsRNA in the central nervous system. One of the additional benefits of using the transgenic UAS/Gal4 system is the ability to fine-tune the degree of Gal4 activity by altering the temperature at which flies are grown [59]. Using this property, we found that HP1a depletion was stronger when larvae were grown at 29°C as opposed to a more mild depletion when larvae were grown at room temperature (measured as 22°C) (Figure S3). When larvae were grown at 29°C, we observed an increase in chromosome segregation errors (Figure S3A-A’), a previously observed phenotype in HP1a mutants [48] and a decrease in survivability (Figure S3B) compared to larvae grown at 22°C.

To determine the role of Aurora B-mediated exclusion of HP1a from acentrics/tethers in nuclear envelope channel formation, we performed live imaging on neuroblasts in which levels of Aurora B and HP1a were modulated, and compared the rates of nuclear envelope channel formation and micronucleation. All larvae were grown in conditions of mild HP1a depletion (22°C). In DMSO-treated neuroblasts (Figure 4A-A’, see Movie S3), we observed that late-segregating acentrics (red arrows) entered daughter nuclei through channels in the nuclear envelope (green arrows) and successfully rejoined the main nuclear mass, consistent with previously reported data [22]. In DMSO + Binucleine-2-treated neuroblasts (Figure 4B-B’, see Movie S4), we observed that late-segregating acentrics (red arrows) failed to form channels in the daughter nuclear envelope and were subsequently locked out of the nuclei to form micronuclei (red arrowheads), as previously reported [22]. In DMSO-treated HP1a RNAi-expressing neuroblasts (Figure 4C-C’, see Movie S5), we observed that late-segregating acentrics (red arrows) entered daughter nuclei through channels in the nuclear envelope (green arrows), successfully rejoining the intact DNA. However, in contrast to the decreased rates of nuclear envelope channel formation and increased micronucleation observed upon inhibition of Aurora B in neuroblasts with wild-type HP1a levels, in DMSO + Binucleine-2-treated HP1a RNAi-expressing neuroblasts (Figure 4D-D’, see Movie S6), we observed that late-segregating acentrics (red arrows) were once again capable of forming nuclear envelope channels (green arrows), through which acentrics rejoined daughter telophase nuclei.

**Figure 4.**
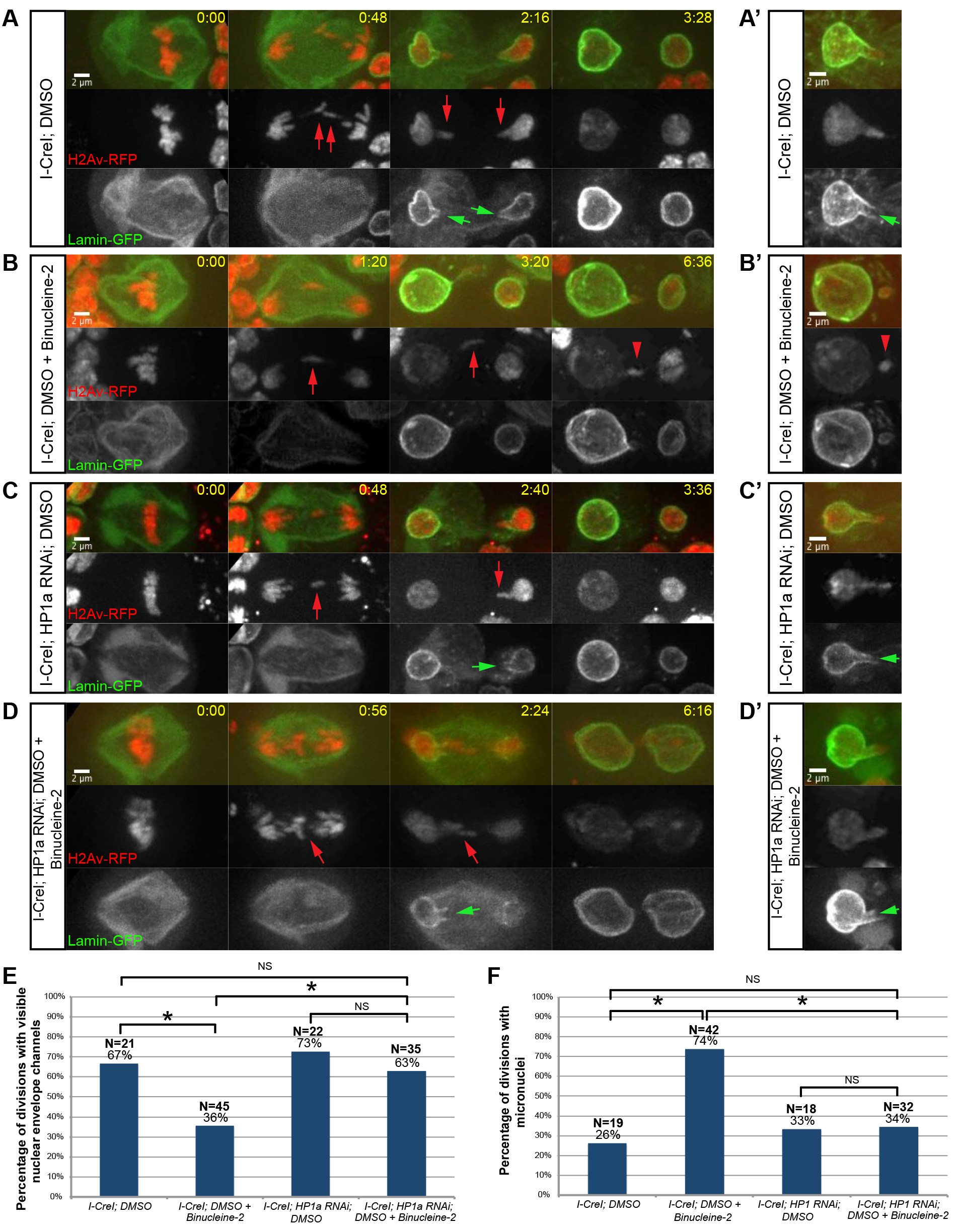
Aurora B-mediated HP1a exclusion from acentrics/tethers results in nuclear envelope channel formation. (A-A’) Stills from time-lapse movies of two different mitotic neuroblasts expressing I-CreI, H2Av-RFP (red), and Lamin-GFP (green) treated with DMSO (see Movie S3), in which acentrics (red arrows) enter daughter telophase nuclei by passing through nuclear envelope channels (green arrows). After acentrics have entered into daughter nuclei, nuclear envelope reassembly completes ((A) 3:28). (B-B’) Stills from time-lapse movies of two different mitotic neuroblasts expressing I-CreI, H2Av-RFP, and Lamin-GFP treated with DMSO + Binucleine-2 (see Movie S4), in which acentrics (red arrows) fail to form nuclear envelope channels, and are instead excluded from the main nuclei as micronuclei (red arrowheads). Micronuclei persist separate from main nuclei well after nuclear envelope reassembly is completed ((B) 6:36). (C-C’) Stills from time-lapse movies of two different mitotic neuroblasts expressing I-CreI, H2Av-RFP, Lamin-GFP, and HP1a RNAi treated with DMSO (see Movie S5), in which acentrics (red arrows) enter telophase daughter nuclei through channels in the nuclear envelope (green arrows). Nuclear envelope reassembly completes after acentrics have rejoined daughter nuclei ((C) 3:36). (D-D’) Stills from time-lapse movies of two different mitotic neuroblasts expressing I-CreI, H2Av-RFP, Lamin-GFP, and HP1a RNAi treated with DMSO + Binucleine-2 (see Movie S6), in which acentrics (red arrows) enter daughter telophase nuclei through channels in the nuclear envelope (green arrows). Acentrics successfully rejoin daughter nuclei by 6:16 (D). (E) Comparison of the percentage of neuroblast divisions in which nuclear envelope channels were observed when I-CreI-expressing neuroblasts were treated with DMSO (67%; N=21) or DMSO + Binucleine-2 (36%; N=45) (asterisk indicates statistically significant to I-CreI; DMSO neuroblasts; p=0.018) and when I-CreI- and HP1a RNAi-expressing neuroblasts were treated with DMSO (73%; N=22) or DMSO + Binucleine-2 (63%; N=35) (asterisk indicates statistically significant to I-CreI; DMSO + Binucleine-2 neuroblasts; p=0.015). NS indicates no statistical significance. (F) Comparison of the percentage of neuroblast divisions in which micronuclei were observed when I-CreI-expressing neuroblasts were treated with DMSO (26%; N=19) or DMSO + Binucleine-2 (74%; N=42) (asterisk indicates statically significant to I-CreI; DMSO neuroblasts; p=0.0005) and I-CreI- and HP1a RNAi-expressing neuroblasts were treated with DMSO (33%; N=18) or DMSO + Binucleine-2 (34%; N=32) (asterisk indicates statistical significance to I-CreI; DMSO + Binucleine-2 neuroblasts; p=0.0006). NS indicates no statistical significance. Time is written as min:sec after anaphase onset. Scale bars are 2 μm. See also Figure S2 and Figure S3.

In total, 67% of DMSO-treated control neuroblasts divisions (N=21) resulted in visible nuclear envelope channels, and inhibition of Aurora B by treating neuroblasts with DMSO + Binucleine-2 resulted in a statistically significant decrease in divisions with visible nuclear envelope channels (36%; N=45) (Figure 4E). In contrast, upon RNAi depletion of HP1a, there was no statistical difference in the percentage of divisions with visible nuclear envelope channels in DMSO-treated neuroblasts (73%; N=22) and DMSO + Binucleine-2-treated neuroblasts (63%; N=35) (Figure 4E). In accord with these observations, we also measured a statistically significant increase in micronucleation from DMSO-treated control neuroblasts (26%; N=19) to DMSO + Binucleine-2-treated neuroblasts (74%; N=42), while depletion of HP1a resulted in no statistical difference between DMSO-treated neuroblasts (33%; N=18) and DMSO + Binucleine-2 treated neuroblasts (34%; N=32) (Figure 4F). Taken together, these results indicate that forming an H3-HP1a complex along the acentric and tether promotes local nuclear envelope assembly, and preventing the formation of this complex, either through Aurora B-dependent H3 phosphorylation or depletion of HP1a, reduces nuclear envelope assembly and facilitates channel formation.

### HP1a exclusion and Aurora B-mediated channel formation are important to prevent micronuclei formation in response to irradiation-induced acentrics

I-CreI produces double-stranded breaks specifically in the centric heterochromatin of the X chromosome and results in acentric fragments of roughly uniform size and heterochromatin content. [20, 25–27]. To test if we would observe similar results regarding HP1a-acentric association and channel formation when acentrics are generated by breaks distributed throughout the genome, we monitored lagging chromosome, HP1a, and nuclear envelope behavior in dividing neuroblasts from larvae exposed to X-irradiation (Figure 5). Ionizing radiation produces single- and double-stranded DNA breaks in both euchromatin and heterochromatin to yield late-segregating acentrics that vary both in size and chromatin composition [60–62]. Despite these differences between I-CreI and irradiation-induced acentrics, in *Drosophila* neuroblasts, acentrics generated by X-irradiation also form BubR1-coated tethers, undergo delayed poleward segregation, and enter daughter telophase nuclei through nuclear envelope channels [18, 22].

**Figure 5.**
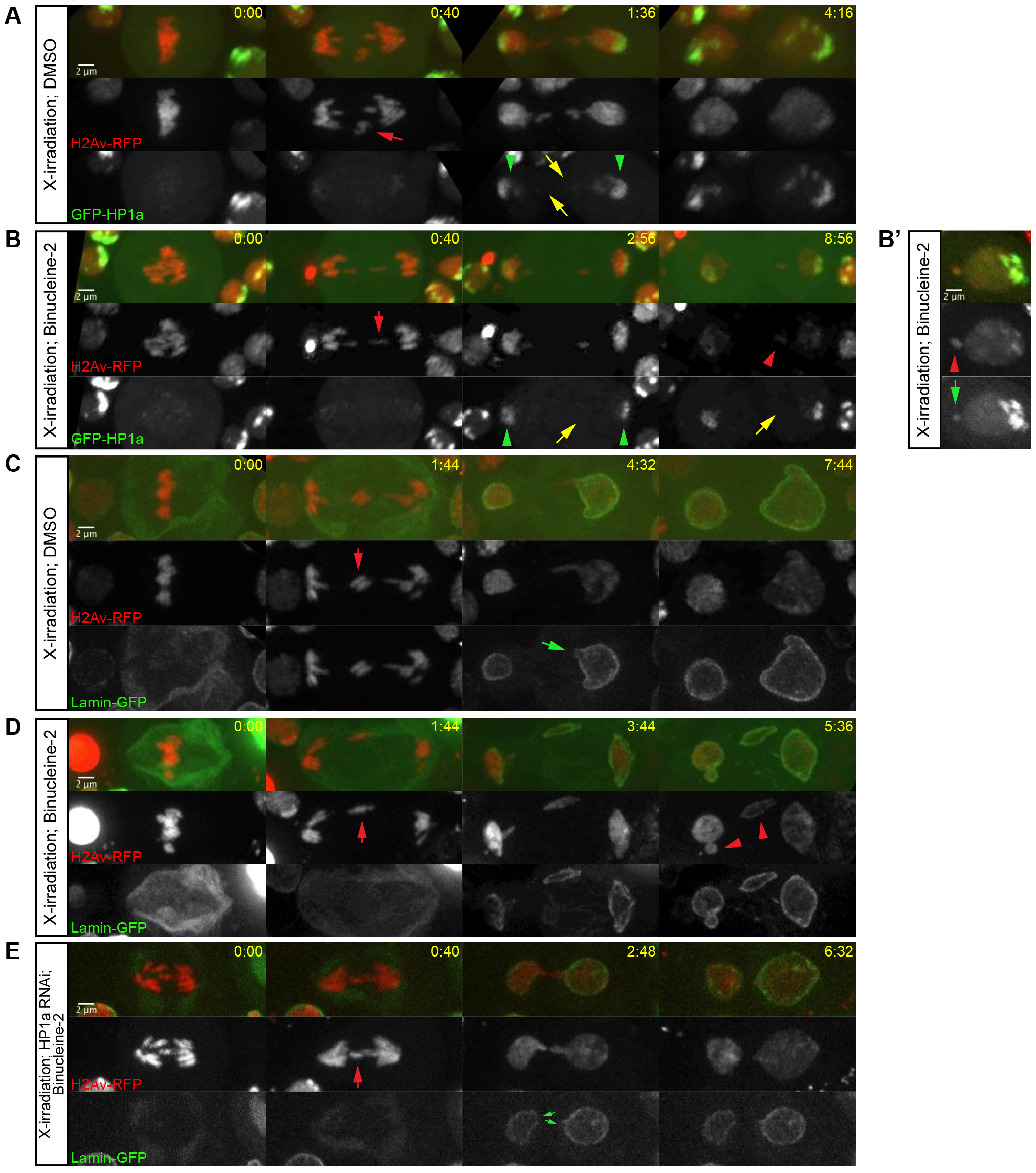
X-irradiation-induced acentrics fail to recruit HP1a and re-enter daughter nuclei through Aurora B-mediated nuclear envelope channels. (A) Stills from a time-lapse movie of an X-irradiated neuroblast expressing H2Av-RFP (red) and GFP-HP1a (green) and treated with DMSO. Acentrics (red arrow) successfully segregate in late anaphase to rejoin daughter nuclei. Despite HP1a association with the main daughter nuclei (green arrowheads), no HP1a was detected on segregating acentrics (yellow arrows). This panel is representative of 5 mitotic divisions observed (5/5 divisions had HP1a recruited to the main nuclei; 0/5 divisions had HP1a recruited to the acentrics; and 4/5 divisions had acentrics that successfully rejoined daughter nuclei). (B-B’) Stills from a time-lapse movie of an X-irradiated neuroblast expressing H2Av-RFP (red) and GFP-HP1a (green) and treated with DMSO + Binucleine-2. (B) Acentric (red arrow) segregates poleward in late anaphase and fails to enter daughter nucleus, forming a micronucleus (red arrowhead). Despite HP1a association with the main daughter nuclei (green arrowheads), no HP1a was detected on the segregating acentric or subsequent micronucleus (yellow arrows). This panel is representative of the 8/11 mitotic divisions observed in which GFP-HP1a was not detected on acentrics. (B’) Shown is a still following mitosis in which after segregating poleward in anaphase, the acentric was coated with GFP-HP1 a and failed to enter the daughter nucleus, instead forming a micronucleus (red arrowhead) coated with GFP-HP1a (green arrow). This panel is representative of the 3/11 divisions which had HP1a recruited to the acentrics. In total 6/9 divisions had acentrics that formed micronuclei. (C) Stills from a time-lapse movie of an X-irradiated neuroblast expressing H2Av-RFP (red) and Lamin-GFP (green) and treated with DMSO. Acentrics (red arrow) successfully segregate in late anaphase and rejoin daughter nuclei by passing through a channel in the nuclear envelope of the daughter nucleus (green arrow). This panel is representative of 6 mitotic divisions observed (4/6 divisions had observable channels; 3/5 divisions had acentrics that successfully rejoined daughter nuclei). (D) Stills from a time-lapse movie of an X-irradiated neuroblast expressing H2Av-RFP (red) and Lamin-GFP (green) and treated with DMSO + Binucleine-2. Acentrics (red arrow) fail to rejoin daughter nuclei as nuclear envelope channels do not form. Instead, acentrics are locked out of the daughter nuclei and form micronuclei (red arrowheads). This panel is representative of 5 mitotic divisions observed (2/5 divisions had observable channels; 4/5 divisions had acentrics that formed micronuclei). (E) Stills from a time-lapse movie of an X-irradiated neuroblast expressing H2Av-RFP (red), Lamin-GFP (Green), and RNAi against HP1a and treated with DMSO + Binucleine-2. Acentrics (red arrow) segregate poleward and rejoin daughter nuclei by passing through channels in the nuclear envelope of the daughter nucleus (green arrows). This panel is representative of 6 mitotic divisions (4/6 divisions had clearly detectable channels; and 0/6 divisions had acentrics that formed micronuclei). Time is written as min:sec after anaphase onset. Scale bars are 2 μm.

To test if Aurora B activity preferentially inhibited HP1a recruitment to X-irradiation-induced acentrics and their tethers, we X-irradiated H2Av-RFP- and GFP-HP1a-expressing neuroblasts. We observed similar patterns of HP1a-acentric dynamics as we did upon I-CreI expression. In control conditions (DMSO-treated neuroblasts), we observed GFP-HP1a exclusion (yellow arrows) from X-irradiation-induced late-segregating chromatin (red arrows) despite its recruitment to the main nuclei (green arrowheads) in 5/5 divisions (Figure 5A), with 0/5 divisions resulting in micronuclei. Upon Aurora B inhibition (DMSO + Binucleine-2-treated neuroblasts), we observed a small increase in GFP-HP1a association (3/11 divisions with detectable GFP-HP1a on acentrics (green arrows)) and a decreased ability for acentrics to rejoin daughter nuclei (6/9 divisions resulting in micronuclei) (red arrowheads) (Figure 5B-B’).

Intriguingly, we observed one way in which the recruitment of HP1a to X-irradiation-induced acentrics differed from that of I-CreI-induced acentrics: whereas I-CreI-induced acentrics recruited HP1a upon Aurora B inhibition in a majority of divisions (55%, N=20) (Figure 3C), X-irradiation-induced acentrics only recruited HP1a upon Aurora B inhibition in a minority of divisions (27%, N=11) (Figure 5B’ green arrow). Nevertheless, these X-irradiation-induced HP1a-free acentrics still formed micronuclei (4/6 observed micronuclei were HP1a-free) (Figure 5B red arrowhead), suggesting that exclusion of HP1a from acentrics is one of several pathways through which Aurora B mediates channel formation and acentric entry into daughter nuclei.

We additionally monitored nuclear envelope channel formation in response to X-irradiation-induced acentrics by X-irradiating H2Av-RFP- and Lamin-GFP-expressing neuroblasts. In X-irradiated DMSO-treated neuroblasts, we observed X-irradiation-induced lagging chromatin (red arrow) reintegrate into telophase daughter nuclei by passing through channels in the nuclear envelope (green arrow) in 4/6 divisions (Figure 5C), with only 2/5 divisions resulting in micronuclei, similar to previous results [22]. Treatment of neuroblasts with DMSO + Binucleine-2 to inhibit Aurora B resulted in decreased channel formation (2/5 divisions with detectable channels) and increased micronucleation (4/5 divisions resulting in micronuclei) (red arrowheads) (Figure 5D). Furthermore, upon reduction of HP1a by RNAi and Aurora B inhibition (DMSO + Binucleine-2-treated neuroblasts), we once again observed increased channel formation (4/6 divisions with detectable channels) (green arrows) and reduced micronucleation (0/6 divisions resulting in micronuclei), suggesting that Aurora B-mediated HP1a exclusion from lagging chromatin to promote channel formation is not simply limited to I-CreI-induced acentrics.

### HP1a specifies preference for nuclear envelope reassembly initiation on the leading edge of segregating chromosomes of the self-renewing neuroblast daughter nucleus

Given that the association of HP1a on acentrics influences local nuclear envelope reassembly at the site of nuclear envelope channels, we hypothesized that HP1a might also play a direct role in global nuclear envelope reassembly in wild-type *Drosophila* neuroblast divisions. Support for this idea comes from studies demonstrating a requirement for HP1α/HP1a to tether heterochromatin to the nuclear envelope following mitosis [52] and in which the expression of a truncated form of HP1 disrupts artificial nuclear envelope assembly in mammalian cells [45], leading to speculation that HP1a may promote nuclear envelope reassembly *in vivo* [63].

To test this hypothesis, we performed live analyses of neuroblast divisions and found GFP-HP1a was always recruited to segregating chromosomes before initiation of nuclear envelope reassembly in the daughter neuroblast (Figure S4A-A’). This is consistent with previous reports of HP1α/HP1a behavior in mitosis [51-52]. On average, we observed that GFP-HP1a was recruited to segregating chromosomes 150 +/− 50 sec (N=16) after anaphase onset and 210 +/− 100 sec (N=15) before nuclear envelope reassembly (Figure S4B). Furthermore, we observed that GFP-HP1a was recruited to the leading edge of segregating chromosomes (Figure S4C). In *Drosophila* neuroblast divisions, which give rise to a self-renewing neuroblast daughter and a ganglion mother cell daughter (GMC), nuclear envelope reassembly initiates on the pole-proximal side of chromosomes segregating to the neuroblast daughter [22, 40]. Therefore, HP1a is located at the proper place and time to mediate nuclear envelope reassembly in neuroblast daughters.

To test the role of HP1a in global nuclear envelope reassembly, we performed live imaging of mitotic neuroblasts expressing H2Av-RFP and Lamin-GFP and monitored nuclear envelope reassembly of neuroblast daughters in wild-type conditions or conditions in which HP1a was strongly depleted through RNAi (Figure 6). As previously reported, in wild-type divisions, we observed a dramatic asymmetry in nuclear envelope reassembly initiation on the self-renewing neuroblast daughter cell [40]. Nuclear envelope reassembly first initiated on the leading, pole-proximal edge of chromosomes segregating to the neuroblast daughter (Figure 6A, green arrows, see Movie S7) before completion on the midzone-proximal face of the segregated chromosomes. As this asymmetry was not as distinct in the differentiating GMC, we focused our studies on the self-renewing neuroblast daughter.

**Figure 6.**
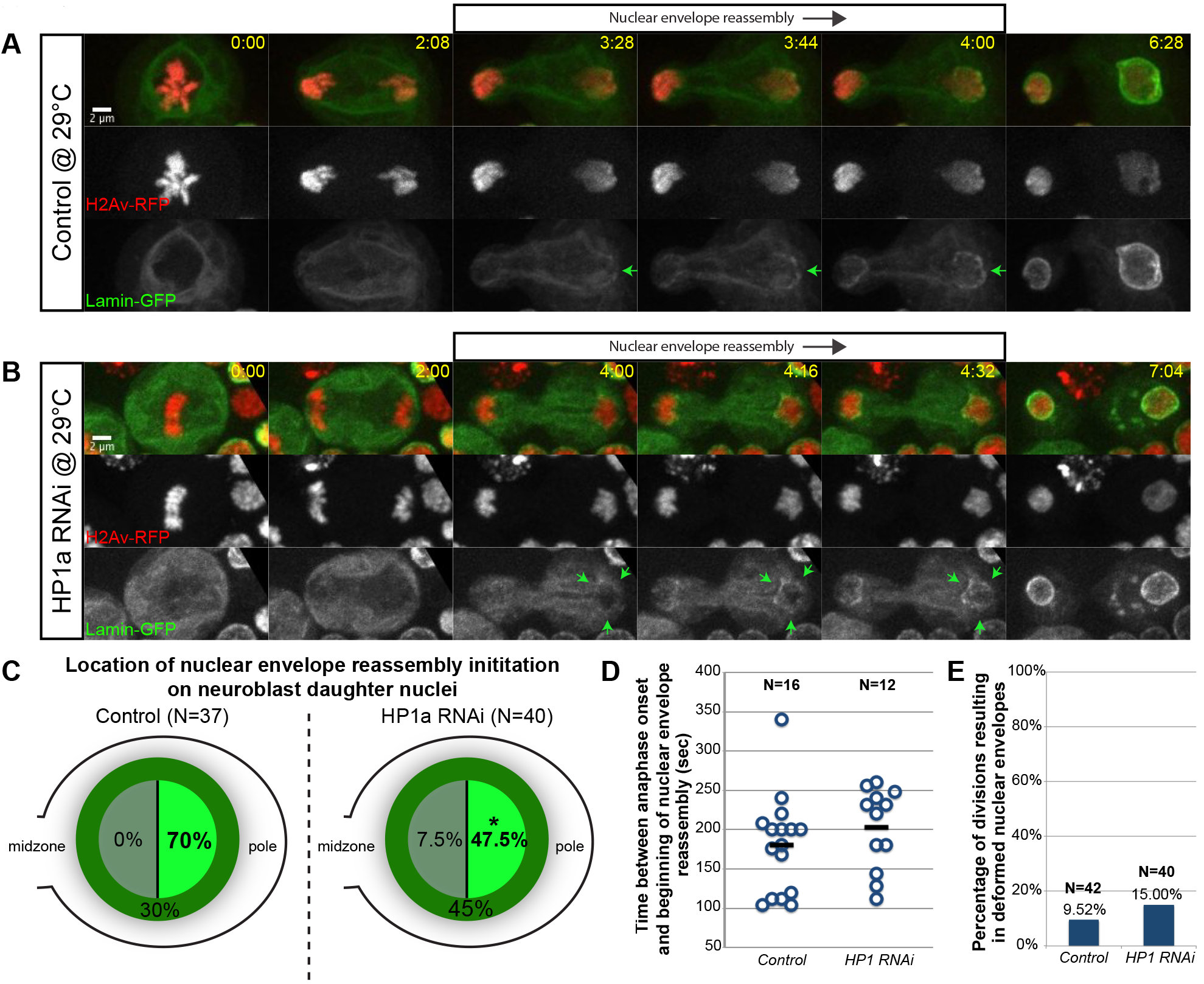
Preferential initiation of nuclear envelope reassembly at the poleward face of segregating chromosomes requires HP1a. (A) Stills from a time-lapse movie of a mitotic neuroblast expressing H2Av-RFP (red) and Lamin-GFP (green) (see Movie S7). In the neuroblast daughter, nuclear envelope reassembly initiates on the pole-proximal face of the segregating chromosomes (green arrows) and wraps around the chromosome mass to completely surround the nucleus. (B) Stills from a time-lapse movie of a mitotic neuroblast expressing H2Av-RFP (red), Lamin-GFP (green), and RNAi against HP1a (see Movie S8). In the neuroblast daughter, nuclear envelope reassembly begins on several sides of the segregating chromosome mass at once (green arrows), including the midzone-proximal face. Nuclear envelope reassembly continues until the nuclear envelope completely surrounds the daughter nucleus. (C) Graph of location frequency for nuclear envelope reassembly initiation in wild-type (left) and HP1a RNAi (right) daughter neuroblasts. Initiation of nuclear envelope assembly on the pole-proximal face of segregating chromosomes in wild-type and HP1a-depleted daughter neuroblasts occurs at a rate of 70% (N=37) and 47.5% (N=40) respectively. Asterisk indicates statistical significance (*p*=0.043). Additionally, another 45% (N=40) of telophase chromosomes initiate nuclear envelope reassembly simultaneously around multiple pole-proximal and pole-distal sites in HP1a RNAi-expressing daughter neuroblasts (compare to 30% (N=37) in wild-type daughter neuroblasts). (D) There is no statistical significance in the time interval between anaphase onset and initiation of nuclear envelope reassembly in wild-type and HP1a-depleted neuroblast divisions. Each circle represents one division. (E) There is no statistical significance in the percentage of divisions resulting in nuclear envelope deformities in wild-type and HP1a-depleted divisions. Time is written as min:sec after anaphase onset. Scale bars are 2 μm. See also Figure S3.

In wild-type neuroblast daughters imaged, we observed nuclear envelope reassembly initiate on the pole-proximal edge of chromosomes 70% of the time, on the midzone-proximal edge of chromosomes 0% of the time, and around all sides of the chromosomes at once 30% of the time (N=37) (Figure 6C). In contrast, in HP1a-depleted neuroblast daughters (Figure 6B, see Movie S8), we observed nuclear envelope reassembly initiate on the pole-proximal edge of chromosomes only 47.5% of the time, on the midzone-proximal edge of chromosomes 7.5% of the time, and around all sides of the chromosome at once (Figure 6B, green arrows) 45% of the time (N=40) (Figure 6C).

In spite of this difference in nuclear envelope reassembly initiation preference, we observed no change in the timing of nuclear envelope reassembly initiation (Figure 6D) or the amount of nuclear envelope deformities (Figure 6E) upon HP1a depletion. Taken together, these results suggest that HP1a specifies the pole-proximal location of nuclear envelope reassembly initiation in self-renewing neuroblast daughter cells.

## DISCUSSION

Despite the inability of acentric chromosomes to form kinetochore-microtubule attachments, studies show that while some acentrics fail to segregate properly [8], others are capable of efficient poleward segregation [18, 24, 62, 64–67]. Failure of acentrics to reincorporate into daughter telophase nuclei leads to the formation of micronuclei, which can cause aneuploidy or DNA damage, and are a hallmark of cancer [9–14]. Recent research has shown that in *Drosophila,* I-CreI-induced acentrics avoid this fate by passing through Aurora B-mediated channels in the nuclear envelope [22].

Here, we found that late-segregating acentrics in anaphase and telophase neuroblast divisions are marked with strong phospho-H3(S10) signal despite the removal of this mark from the main nuclei (Figure 2). The strength of this signal is dependent upon Aurora B (Figure S1). Interestingly, during late anaphase when the normal chromosomes begin to reassemble a nuclear envelope, the acentrics remain nuclear envelope-free [22], suggesting an inverse correlation between nuclear envelope reassembly and phospho-H3(S10) modification. This finding is consistent with studies suggesting that, in addition to the phosphorylation state of nuclear envelope components [68–69], nuclear envelope reassembly is also regulated by various chromatin remodeling events, including the removal of phospho-H3(S10) [70–71].

Our studies also demonstrate that HP1a is excluded from late-segregating I-CreI-induced acentrics, despite HP1a recruitment to the main nuclei (Figure 3). We note the chromatin makeup of I-CreI-induced acentrics should be sufficient to recruit HP1a, as I-CreI creates double-stranded breaks in the pericentric region of the X chromosome, resulting in an acentric fragment which should contain a large portion of heterochromatin [20, 25]. Presumably, the difference in recruitment of HP1a to acentrics and the main nuclei is due to the difference in the phosphorylation state of H3(S10) on acentrics and the main nuclei, consistent with the view that Aurora B-mediated phospho-H3(S10) is prohibitive to HP1α/HP1a binding [47–48]. Indeed, in support of this view, inhibition of Aurora B kinase activity resulted in increased HP1a association with acentrics (Figure 3).

Furthermore, we find that acentrics possessing high levels of HP1a were largely unable to reintegrate into daughter telophase nuclei and instead formed micronuclei (Figure 3). Under the same Aurora B inhibition conditions in which we detected HP1a association with acentrics, we observed increased nuclear envelope reassembly around acentrics and decreased nuclear envelope channel formation on main nuclei (Figure 3-4). We therefore proposed a pathway in which Aurora B activity mediates nuclear envelope channel formation by blocking anaphase HP1a recruitment to acentrics and their associated tethers (Figure 1). This model predicts that 1) in wild-type conditions, Aurora B activity would inhibit formation of an H3-HP1a complex on the acentric and tether and lead to slow nuclear envelope assembly at these sites and the formation of a channel; 2) when Aurora B is inhibited, H3 on the acentric and tether would bind to HP1a which would stimulate nuclear envelope assembly at these sites and prevent channel formation; and 3) when Aurora B is inhibited and HP1a is depleted, no H3-HP1a complex would form on the acentric and tether, leading to slow nuclear envelope assembly at these sites and the formation of a channel, reminiscent of wild-type conditions (Figure S2). Our data show upon co-depletion of HP1a with Aurora B inhibition, nuclear envelope reassembly on acentrics is reduced and channel formation occurs at frequencies similar to those detected in wild-type Aurora B and HP1a conditions (Figure 4). Essentially, depletion of HP1a masks the phenotype of Aurora B inhibition, evocative of a classic epistatic relationship, in which Aurora B mediates nuclear envelope channel formation through preferentially excluding HP1a from acentrics and their tethers.

Intriguingly, channel formation mediated by HP1a exclusion is similar to the mechanism of human polyomavirus egress from its host nucleus. Viral angoprotein binds to HP1α/HP1a and disrupts its binding with the inner nuclear membrane protein LBR, causing sections of weakened nuclear envelope through which virions leave the nucleus [72]. Thus regulation of chromatin-HP1 a-nuclear envelope interactions may represent a conserved method for bypassing the barrier of the nuclear envelope.

Significantly, by generating acentrics through X-irradiation, we found that the general pattern of HP1a recruitment driving nuclear envelope reassembly on acentrics was not dependent upon the system used to generate acentrics or due to the exact physical nature of an I-CreI-induced acentric (Figure 5). However, we note that while micronuclei derived from I-CreI-induced acentrics were generally HP1a-coated, a higher proportion of micronuclei derived from irradiation-induced acentrics were HP1a-free. It is tempting to speculate that these HP1a-free micronuclei were derived from largely euchromatic acentrics that simply lack a sufficient amount of heterochromatin to recruit HP1a. This observation suggests that Aurora B may mediate acentric entry into daughter nuclei through multiple pathways of which preferential exclusion of HP1a is one. It is possible these HP1a-free acentrics remain capable of recruiting a nuclear envelope through an HP1a-independent pathway, perhaps involving an interaction between LEM domain-containing inner nuclear membrane proteins and the DNA-crosslinking factor barrier-to-autointegration factor [73–74].

To support our finding that HP1a promotes nuclear envelope assembly, we examined its role in assembling the nuclear envelope on normal intact chromatin. We observed HP1a localize to the leading edge of segregating chromosomes before nuclear envelope reassembly (Figure 5). As previously reported, we observed nuclear envelope reassembly initiate on the leading edge of chromosomes segregating to daughter neuroblasts, proceeding to wrap around and complete reassembly on the midzone-proximal face of the nascent nucleus [22, 40, 75]. However, reducing HP1a levels disrupts the preferential nuclear envelope assembly on the pole-proximal face of the segregating chromosomes (Control = 70% pole-proximal initiation; HP1a depletion = 47.5% pole-proximal initiation) (Figure 6). This result is in accord with a growing body of evidence suggesting HP1 proteins may play key roles in nuclear envelope reassembly. For example, in mammalian cells, HP1α/HP1a recruits PRR14 to segregating chromosomes where it tethers heterochromatin to the nuclear envelope [52], and *in vitro* experimentation shows HP1β/HP1b to be important for recruiting nuclear envelope components to interphase-like chromatin [15]. In addition, depletion of the PP1γ subunit Repo-Man leads to retained phospho-H3(S10) marks on chromatin, loss of HP1α/HP1a recruitment to mitotic chromosomes, and defects in nuclear envelope reassembly [70]. One mechanism by which HP1a might bias nuclear envelope reassembly to initiate on the leading edge of segregating chromosomes is by enhancing the natural ability of nuclear envelope components to bind to chromatin [76].

Despite the clear preference for initiation of nuclear envelope reassembly on the leading edge of chromosomes segregating to daughter neuroblasts, we observed no such preference on the chromosomes segregating to the daughter GMC (Figure 6). It is possible this difference is due to the relatively small size of the GMC daughter, or that in *Drosophila* neuroblast divisions, the endoplasmic reticulum, from which the nuclear envelope extends during mitotic exit, is asymmetrically localized to the spindle pole of the neuroblast daughter [77].

Our data also address a key question regarding nuclear envelope channel formation: how do late-segregating acentrics near the spindle midzone act at a distance to influence nuclear envelope reassembly dynamics on main nuclei near the poles? In our system, we believe there are two pools of Aurora B: a constitutive midzone-based pool [30, 54], and a tether-based pool, which stretches from the acentric to the main nucleus [18]. Given that nuclear envelope channels are only observed on the main nuclei when acentrics and the tether-based pool of Aurora B are present, we proposed that the pool of Aurora B responsible for channel formation is the tether-based Aurora B [22]. Our observation of phospho-H3(S10) “hotspots” on the main nuclei at sites closest to acentrics is consistent with the hypothesis that tether-based Aurora B activity controls channel formation (Figure 2A, yellow arrowheads). Since the midzone pool of Aurora B is confined away from the main telophase nuclei, it is probable that these hotspots are due to the activity of Aurora B along tethers, which stretch from acentrics and contact main nuclei at sites closest to the acentrics. These phospho-H3(S10) hotspots could then locally prevent HP1a association and nuclear envelope reassembly. Thus, while nuclear envelope reassembly can still proceed around the rest of the nucleus, it is inhibited at the site of these hotspots, resulting in channels.

In summary, our results reveal a novel mechanism by which genome integrity is maintained. Late-segregating acentric fragments pose a significant hazard, as they are at high risk of forming micronuclei that induce dramatic rearrangements in the genome [13–14, 29]. Consequently it is likely that cells have evolved mechanisms to prevent the formation of micronuclei. Here, we provide evidence for one such mechanism in which Aurora B-mediated inhibition HP1a-chromatin association during anaphase/telophase prevents the formation of micronuclei from late-segregating acentric fragments.

## METHODS

### Fly stocks

All stocks were raised on standard *Drosophila* food [78]. Chromosome dynamics were monitored using H2Av-RFP (stock #23651, Bloomington *Drosophila* Stock Center [BDSC], Bloomington, IN). The following Gal4 drivers were used: elav-Gal4 [79], Wor-Gal4 [80], and Actin-Gal4 (#25708, BDSC; [81]). To monitor nuclear envelope dynamics, we expressed UAS-lamin-GFP (#7376, BDSC) driven by elav-Gal4. HP1a localization was assessed through use of GFP-HP1a (#30561, BDSC). UAS-ial-dsRNA (#28691, BDSC) driven by Wor-Gal4 was used to deplete Aurora B. UAS-Su(Var)205-dsRNA (#33400) driven by either Actin-Gal4 or elav-Gal4, depending on the experiment, was used to deplete HP1a.

### Fixed neuroblast cytology

Crawling female 3^rd^ instar larvae bearing either hs-I-CreI and Wor-Gal4 or hs-I-CreI, Wor-Gal4, and UAS-ial-dsRNA were heat shocked for 1 hour at 37°C. Following 1 hour recovery at room temperature, brains were dissected in 0.7% NaCl then fixed in 3.7% formaldehyde for 30 min. Brains were washed in 45% acetic acid in PBS for 3 min then placed between siliconized coverslips and glass slides in 60% acetic acid in PBS. Brains were squashed by tracing over coverslips with watercolor paper. Slides were frozen in liquid nitrogen for 10 min and then washed in 20% ethanol for 10 min at −20°C. After washing with PBST (10 min) and PBS (2 × 5 min), slides were blocked in a 5% dried milk solution in PBST for 1 hour. Samples were incubated with rabbit anti-histone phospho-H3(S10) antibody (abcam #ab5176) at a 1:500 dilution overnight at 4°C. Samples were subsequently washed in PBST before incubation with goat anti-rabbit-alexa488 (ThermoFisher #A-11008) at a 1:300 dilution for 1 hour at room temperature. Slides were washed in PBST, counterstained with DAPI in vectashield, and imaged the following day. Procedure adapted from [82–83].

### Quantitative immunofluorescence imaging

Fixed slides were imaged on a Leica DMI6000B wide-field inverted microscope equipped with a Hamamatsu EM CCD camera (ORCA C9100-02) with a binning of 1 and a 100x Plan-Apochromat objective with NA 1.4. For experiments determining the ratio of phospho-H3(S10)/DNA on the acentrics vs. on the main nuclei, phospho-H3(S10)/chromatin pixel intensity was determined in ImageJ (National Institutes of Health, Bethesda, MD) by drawing individual region of interests (ROIs) around both main nuclei and the acentrics (as determined from DAPI staining) from sum projections of all z-slices in which the nuclei and acentrics were in focus. Corrected total fluorescence (CTF) was calculated for each ROI (acentrics and nuclei) for both DAPI and phospho-H3(S10) channels by subtracting the product of the ROI area and the mean pixel intensity of an arbitrarily-selected background region from the measured integrated density of the ROI. For each division set, CTFs were averaged for the two main nuclei and for the acentrics when more than one acentric ROI was drawn. phospho-H3(S10)/DAPI ratios were calculated by dividing the CTF of phospho-H3(S10) by the CTF of DAPI for the averaged acentrics and the averaged nuclei. To determine the fold change for acentrics vs. main nuclei phospho-H3(S10)/DAPI ratios, the phospho-H3(S10)/DAPI ratio of acentrics was divided by that of the main nuclei for each imaged division.

To compare phospho-H3(S10) levels on acentrics in I-CreI vs. I-CreI; Aurora B RNAi neuroblasts, control and Aurora B-depleted brains were imaged at the same laser settings. Quantification of phospho-H3(S10)/DAPI ratios were calculated as detailed above.

### Live neuroblast cytology

For experiments involving acentrics, crawling female 3^rd^ instar larvae bearing hs-I-CreI, elav-Gal4, H2Av-RFP, and a combination of GFP-HP1, UAS-Lamin-GFP, and/or UAS-Su(Var)205-dsRNA were heat shocked for 1 hour at 37°C. Larvae were allowed to recover for at least 1 hour following heat shock. For experiments with no acentrics, female 3^rd^ instar larvae bearing elav-Gal4, H2Av-RFP, and a combination of GFP-HP1, UAS-Lamin-GFP, and/or UAS-Su(Var)205-dsRNA were used. Brains were dissected in PBS and gently squashed between a slide and coverslip [84]. Neuroblasts along the periphery of the squashed brain provided the best imaging samples. Slides were imaged for up to 1 hour.

Data from time-lapse imaging experiments were acquired with both a Leica DMI6000B wide-field inverted microscope equipped with a Hamamatsu EM CCD camera (ORCA C9100-02) with a binning of 1 and a 100x Plan-Apochromat objective with NA 1.4 and an inverted Nikon Eclipse TE2000-E spinning disk (CSLI-X1) confocal microscope equipped with a Hamamatsu EM-CCD camera (ImageE MX2) with a 100X 1.4 NA oil immersion objective. Successive time points were filmed at 20 sec for the wide-field microscope and 8 sec for the spinning disk microscope. Spinning disk images were acquired with MicroManager 1.4 software.

### Small molecule inhibition of Aurora B kinase

For experiments involving the depletion of Aurora B kinase, following dissection, brains were washed in a 25.5μM solution containing Binucleine-2 (Sigma B1186) for 5 min, after which brains were squashed in PBS between a slide and coverslip. For control experiments, dissected brains were washed in 0.15% DMSO (final concentration of DMSO in solution used to dissolve Binucleine-2) for 5 min and then squashed in PBS between a slide and coverslip. Neuroblasts entering anaphase were selected for imaging, and slides were imaged for only one division.

### Temperature-regulated expression of RNAi and lethality studies

Flies bearing either Actin-Gal4 or Actin-Gal4 and UAS-Su(Var)205-dsRNA were grown at room temperature (measured as 22°C) until they reached 3^rd^ instar stage. At this point, larvae were collected into vials and either allowed to continue to grow at room temperature or were shifted to grow at 29°C. Survivability was determined by counting the number of adult flies that eclosed in each vial.

### Irradiation studies

Crawling 3^rd^ instar larvae were placed in an empty plastic vial and irradiated with 604.8 rads using a Faxitron CP160 X-ray machine. Larvae were allowed to recover for at least 1 hour before brains were dissected and mitotic neuroblasts were imaged as described above.

### Statistical analyses

Statistical analyses were determined by chi-square tests (Figure 3C-D, Figure 4E-F, Figure 6C and E) and t-tests (Figure 6D and Figure S3B) performed in R (R Core Team (2014)).

### Figure preparation

Figures were assembled using ImageJ software and Adobe Illustrator (Adobe, San Jose, CA). Graphs were assembled in Microsoft Excel (Microsoft, Redmond, WA). Selected stills from experiments involving live imaging were adjusted for brightness and contrast using ImageJ to improve clarity.

## AUTHOR CONTRIBUTIONS

B.W. and W.S. conceived experiments and wrote the manuscript. B.W. performed experiments.

## ACKNOWLEDGEMENTS

We thank W. Saxton and S. Strome for use of their equipment and B. Abrams for additional help with microscopy. We thank the members of the C. Forsberg Lab for their assistance with irradiation experiments. We also thank G. Karpen and P. O’Farrell for their helpful advice. This work was funded by the National Institute of Health R01- GM120321 grant awarded to W.S.; B.W. was also supported by National Institute of Health grant 5T32GM008646-18. The authors declare no competing interests.

